# Adaptive Resilience: Agarikon Mycelium Modulates Immune Responses and Provides Oxidative Stress Buffering in Human Immune Cells

**DOI:** 10.64898/2026.07.10.737773

**Authors:** Elizabeth Doar, Jessica Kishiyama, Zolton J. Bair, Paul Stamets, Chase Beathard

**Affiliations:** Department of Research and Development, Fungi Perfecti, LLC, Olympia, WA, USA

**Keywords:** *Fomitopsis officinalis*, *Laricifomes officinalis*, agarikon, mycelium, immune modulation, antioxidant, cytokine signaling

## Abstract

Agarikon (*Fomitopsis officinalis*, syn. *Laricifomes officinalis*) is a fungus with millennia of traditional use across many cultures with modern research supporting the antimicrobial, antiviral, anticancer, antioxidant, and immune-modulating properties of both mycelium and fruit body. Due to the slow growth and old-growth forest habitat of agarikon fruit bodies, its mycelium represents an easily cultivated immunomodulating and stress buffering preparation, underscored by recent clinical trials of a blend of agarikon and *Trametes versicolor* mycelium. We investigated the transcriptomic effects of agarikon mycelium in human peripheral blood mononuclear cells (PBMCs) under both basal and LPS-stimulated conditions, alongside evaluations of antioxidant, iron chelating, and kinase binding activity. Two agarikon fractions were also assessed for effects on cell viability and proliferation and induction of select cytokine targets. Under basal conditions, agarikon mycelium selectively engaged the innate immune response through IL-1 and NF-κB axes, balanced by increases in anti-inflammatory mediators such as IL-1RA and decreases in toll-like receptor transcripts. Under LPS-stimulated conditions, this innate immune response was modified, with measured increases in immune effectors (including *TLR5*) observed in response to induced stress, alongside accompanying transcript decreases in cytokine pathways that can overstimulate the immune system. Overall, agarikon mycelium demonstrated a coordinated, context-dependent immune response profile, supporting its stress-buffering and immune-modulating potential and warranting continued clinical validation.

## 1 Introduction

Agarikon (*Fomitopsis officinalis*, syn. *Laricifomes officinalis*) is a medicinal mushroom with millennia of traditional use in many cultures, with preparations included in ancient pharmacopoeias such as Dioscorides’ De Materia Medica (1, 2, 3). Modern research has supported the antimicrobial, antiviral, anticancer, immunomodulatory, and antioxidant properties of individual components and whole extracts of agarikon mycelium (2, 4, 5, 6, 7, 8, 9, 10, 11, 12) and fruit body (4, 13, 14, 15). The antioxidant and anticancer effects of agarikon treatment are partially enacted by its distinct immunomodulatory capacities and mediated by the nuclear factor kappa-light-chain-enhancer of activated B cells (NF-κB) pathway among others, as demonstrated *in vitro* using cancer cell lines and *in vivo* using zebrafish models (13, 14, 15).

Fungi can produce compounds unique to developmental stage, substrate, and extract type, which influence the immunological effects of fungal treatments (16, 17). For instance, two chlorinated coumarins have been isolated from agarikon mycelium, with structure-dependent activity against *Mycobacterium tuberculosis*, expanding the reported bioactive metabolite space of this species beyond its well-studied terpenoid fraction (6). Additionally, investigations of the antiviral activity of agarikon have thus far centered on the action of mycelial fractions, in particular against pox viruses (12) and influenza A (18). A specific agarikon fruit body fraction was found to act as a toll-like receptor (TLR) agonist by activating NF-κB and the innate immune system via TLR2 and TLR4, and bolstered the adaptive immune system by binding to and preventing the interaction of programmed cell death protein 1 (PD-1) and its ligand 1 (PD-L1) and vascular endothelial growth factor (VEGF) and its receptor (VEGFR-2), which inhibits tumor angiogenesis (14, 15). A broader fraction of agarikon fruit body was also observed to engage this pathway by upregulating NF-κB inhibitor alpha (*NFKBIA*), NF-κB subunit 1 (*NFKB1*), and transcription factor p65 (*RELA*) protein products in HepG2 cancer cells as part of its anticancer effect (13). However, some preclinical studies suggest that high concentration polysaccharide extracts from fruit bodies of certain species, including *Ganoderma lucidum* and *Hericium erinaceus*, may reduce viability (19) or induce stress responses (17) in human immune cells. Collectively, these findings suggest that immunological activity elicited by fungal-derived preparations is highly dependent on biological source and extraction method, necessitating functional evaluation across tissue types and solvent-based extracts to characterize context-dependent immune responses *in vitro*.

Because agarikon fruit bodies are slow growing and largely limited to old-growth forest habitats, contributing to its Endangered status on the IUCN Red List of Threatened Species, aseptically cultured mycelium has emerged as a controlled and scalable method for accessing its bioactive potential without reliance on wild harvest (11, 20). Accordingly, one cryopreservation program has established a repository containing at least 87 genetically distinct agarikon strains, supporting long-term conservation of its genetic diversity (21). Recently, an annotated nuclear genome of agarikon has been made publicly available for the first time, providing foundational resources for understanding its genetic architecture and biosynthetic potential, and reflecting its status as a phylogenetically distinct polypore and sole member of the family *Laricifomitaceae* (22). Building on the genetic, ecological, and functional rationale for investigating agarikon mycelium, this study evaluated its potential as a cytocompatible and effective immune- and oxidative stress-modulating treatment in human cells to assess its suitability for medicinal use.

## 2 Materials and Methods

### 2.1 Sample Preparation

Two different extracts (hydroethanolic, EtOH and ethyl acetate, EtOAc) of agarikon mycelial samples were prepared. EtOH extracts were made by combining 20 g of Host Defense agarikon powder (*F. officinalis* mushroom mycelium and fermented brown rice biomass) with 50 g 95% ethanol and 50 g ultrapure water. The sample was macerated for 120 h at room temperature, centrifuged at 600 x g for 10 min, and the supernatant decanted. Centrifugation was repeated until the pellet was nearly dry. After maximum supernatant recovery, samples were rotary evaporated to reduce fluid volume and frozen at –80 °C. Samples were then lyophilized to dryness, transferred to a new sterile tube, and stored at room temperature. Then, samples were reconstituted in water, DMSO, or PBS, passed through a 0.2 μm polypropylene filter, and used for functional assays. EtOAc samples were prepared by milling lyophilized agarikon mycelium to reduce particle size. 2 g of milled agarikon powder was combined with 10 mL of ethyl acetate, sonicated for 10 min, vortexed for 30 min, and set to macerate for 24 h at room temperature. After maceration, the sample was centrifuged at 600 x g for 10 min, and the supernatant decanted. Centrifugation was repeated until the pellet was nearly dry. After maximum supernatant recovery, the samples were rotary evaporated to remove any excess solvents and lyophilized to dryness. Samples were reconstituted in DMSO and passed through a 0.2 μm polypropylene filter before being used for functional assays.

### 2.2 Cell Viability and Proliferation Testing

#### 2.2.1 MTT Evaluation of Cell Viability and Cytotoxicity

Before culturing PBMCs for RNA extraction and additional assays, MTT assays (ATCC, 30-1010K) were performed to evaluate cell viability (as based on metabolic activity) following treatment with agarikon extract preparations. MTT assays were performed in accordance with the manufacturer’s kit instructions, but with a decrease in volume from 110 μl to 55 μl reactions to conduct the assay in a 384-well plate, with a minimum of n = 4 replicate wells per treatment concentration. Standard human PBMCs (ATCC, PCS-800-011) were thawed, centrifuged, and washed with 37 °C RPMI 1640 media to remove DMSO used for cryopreservation purposes. Cells were incubated in 15 mL of warmed RPMI 1640 (10% FBS, heat inactivated, Sigma-Aldrich, F1051-100ML) overnight in a T75 flask at 37 °C in a 5% CO_2_ atmosphere to recover from the freeze/thaw process. Cell counts were determined via hemocytometer after staining with 0.4% trypan blue diluted in PBS. Following a 24 h growth phase within the flask, cells were transferred to a 96-well plate (2 × 10⁶ cells per well).

Treatments were introduced approximately 24 h after seeding, and plates were incubated for 18-24 h post-treatment before addition of the MTT reagent and further incubating at 37 °C in a 5% CO_2_ atmosphere for 4-24 h to facilitate formazan crystal formation. Detergent was then added for 4-24 h until crystals were completely solubilized. Absorbance readings were recorded at 570 and 660 nm, and sample values were calculated by subtracting values at 660 nm from values at 570 nm. Percent cell viability was evaluated relative to either the PBS or DMSO vehicle controls, according to each sample’s vehicle solvent.

#### 2.2.2 Fluorometric Evaluation of Cell Proliferation

Cells were also evaluated for cellular proliferation levels under both basal and LPS inflammatory conditions via fluorometric assessment (Invitrogen, C35006) to more thoroughly explore the cytocompatibility of agarikon mycelium carried in a variety of solvents in human cell culture. Standard human PBMCs were cultured and treated as described for MTT assay and then centrifuged at 100 x *g* for 6 min. Cell media was aspirated and frozen at –20 °C for use in ELISA quantification. A working solution of 1X HBSS buffer containing CyQUANT NF dye reagent (1:500 dilution) was prepared in accordance with the nonadherent cell protocol provided. Cells were washed with PBS and aspirated before 100 µL of combined 1X HBSS buffer and dye reagent was added to each well. Plates were incubated at 37 °C for 60 min before being read fluorometrically (485 nm excitation, 528 emission), and cell proliferation values were calculated as percentages relative to the DMSO vehicle control.

#### 2.2.3 Cell Imaging

To produce representative images of cells for MTT and fluorometric cell proliferation assays, pictures of agarikon- and vehicle-treated cells were obtained either by staining with 0.4% trypan blue diluted in PBS and imaging directly using a Leica DMi1 microscope and Leica MC170 HD digital microscope camera, or by resuspending cells in 1X HBSS buffer and dye binding solution (Invitrogen, C35006) and imaging with the Leica DMi1 and Leica MC170 under UV illumination, adapted from (23). ImageJ (v1.54p) was used for analysis.

### 2.3 PBMC Culture for mRNA-Sequencing

mRNA-sequencing (RNA-Seq) was used to assess global gene expression changes in PBMCs following agarikon treatment under basal and LPS-stimulated conditions. After an initial 24 h incubation in 12-well plates, PBMCs were treated and incubated for a further 24 h, including 100 ng/mL LPS for inflammatory condition samples. For harvest, cells and media were aspirated, pipetted into sterile 1.5 mL microcentrifuge tubes, washed with PBS, and aspirated again before freezing cell pellets in liquid nitrogen prior to storage at –80 °C.

Before starting RNA extraction, surfaces and tools were cleaned with RNaseZap (ThermoFisher, AM9780). Using the RNeasy Mini Kit (Qiagen, 74104), 350 µL of RLT buffer was added to each microcentrifuge tube after flicking to loosen the cell pellet. Cells were lysed by vortexing for 1 min, and the supernatant was transferred to a new sterile 1.5 mL microcentrifuge tube. The manufacturer’s recommended extraction procedure for mammalian cells was followed, including the optional step of placing columns into new collection tubes and centrifuging for 1 min at 13,000 x *g* immediately before elution. 30 µL of RNase-free water was used for elution, and columns were incubated at room temperature for 1 min before proceeding. RNA quality (A260/280 ratios) and concentrations of samples were determined spectrophotometrically prior to DNase digestion, where 30 µL of eluted RNA was added to 3 µL of reaction buffer and 3 µL of DNase I (ThermoFisher, EN0521). Samples were incubated at 37 °C for 30 min before 3 µL of 50 mM EDTA was added, then incubated at 65 °C for 10 min. All samples were stored at –80 °C and shipped to Novogene Corporation Inc. (Sacramento, California) on dry ice for RNA-Seq.

### 2.4 RNA-Seq and Analysis

Extracted samples proceeded to sequencing if they had a total RNA quantity of ≥100 ng and a spectrophotometric A260/280 ratio of 1.95 or greater. Novogene’s mRNA-Seq service was used, performed on an Illumina NovaSeq X-Plus platform with a paired-end read length of 150 bp. At least 20 million read pairs per sample were generated and aligned to the reference genome *Homo sapiens* GRCh38 (hg38); RNA-Seq data are summarized in Supplementary Tables S1 and S2. NovoMagic (Novogene’s RNA-Seq platform) was used to generate differentially expressed gene (DEG) lists and dot plots summarizing KEGG and Reactome pathways. DESeq2 analysis, applying the Benjamini-Hochberg false discovery rate (FDR) correction with an adjusted *p*-value (*p*adj) cutoff of ≤ 0.05, was used to identify significant DEG and pathway results (24). PCAGO was used to assess variability via principal component analysis (PCA), RStudio 2026.05.0 (R 4.6.0) was used to create bar and line graphs, volcano plots, and heatmaps, and KEGG Mapper – Color was used to generate and color code KEGG plots.

### 2.5 Cytokine Activity Assays

Protein-level confirmation of select RNA-Seq outcomes was pursued using ELISAs to provide a complementary evaluation of the functional immune response in PBMCs. Standard human PBMCs were cultured as described for evaluation of cell proliferation and viability, and conditioned cell media was aspirated and used for cytokine analysis via ELISA. Cytokine concentrations were quantified using commercially available ELISA kits for interleukin 8 (IL-8; R&D Systems, DY208-05), interleukin-1 receptor antagonist (IL-1RA; R&D Systems, DY280-05), and interleukin-1 alpha (IL-1α; R&D Systems, DY200-05). All ELISAs were run according to manufacturers’ instructions besides volume modifications to accommodate a 384-well format. IL-8 and IL-1RA used PBS and DMSO as vehicle controls, and IL-1α used LPS as a vehicle control.

### 2.6 MAPK Binding Assays

Investigation of the mitogen-activated protein kinase (MAPK) binding activity of agarikon mycelium was performed using Eurofins’ DiscoverX KINOMEscan profiling service to explore secondary signaling pathways induced by this strain of agarikon that could influence immune responses. An EtOAc extract was selected to enrich for hydrophilic and lipophilic secondary metabolites. c-Jun N-terminal kinase 3 (JNK3) and tropomyosin receptor kinase A (TRKA) were included in KINOMEscan, a competitive binding assay used to evaluate the ability of agarikon mycelium extracts to disrupt interactions between kinases and immobilized ligands. Results are reported as percent control values, where lower values indicate stronger kinase binding interactions relative to the DMSO control (25, 26).

### 2.7 DPPH Assays

DPPH radical scavenging assays were employed to assess general antioxidant potential of agarikon and provide functional corroboration of the stress- and redox-related pathways explored via transcriptomics. A 100 µg/mL DPPH reagent (Sigma-Aldrich, 300267) stock was prepared in 99% ethanol and calibrated by adding 60 µL of DPPH stock to 40 µL of sterile water in a single well of a 384-well plate and ensuring absorbance values fell between 1.0 and 1.1 when read spectrophotometrically at 517 nm. DPPH reagent was included as a maximal signal control, and agarikon treatment was serially diluted in sterile water or 99% ethanol before being added to a 384-well plate at a volume of 75 µL per well. Background was evaluated at 517 nm before adding 25 µL DPPH stock to each well and reading at 517 nm every 5 min for 60 min, shaking between read points. Final absorbance values were obtained by subtracting the average background absorbance at 60 min from all other wells, and percent radical scavenging activity was calculated by subtracting sample absorbance from the maximal signal control, dividing by the maximal signal control, and multiplying by 100, where the maximal signal control was defined as the absorbance of DPPH reagent alone at 60 min.

### 2.8 Evaluation of Ferrous Iron Chelating Ability

In addition to general antioxidant assessment and transcriptomic analysis of iron-related signaling pathways, the iron chelation activity of agarikon mycelial extract was measured to directly quantify its iron-sequestering potential. Following (27) and (28), ferrous iron (Fe²⁺) chelating activity was measured by quantifying the reduction in color intensity after inhibition of the purple ferrozine–Fe²⁺ complex. Water was used for all sample and component dilutions and controls, and an EDTA serial dilution (100 µM-0.78 µM) was included to express iron chelation ability in EDTA µM equivalents. In 1.5 mL microcentrifuge tubes, 500 µL of control, blank, or sample was added to 900 µL of sterile water, and 100 µL of 0.6 mM FeCl_2_ (ThermoFisher, 389350250) was added to all reactions except the blank, where it was replaced with 100 µL water. Reaction tubes were vortexed, incubated for 10 min at room temperature with additional vortexing every 2 min, and then 100 µL 5 mM ferrozine (ThermoFisher, AC410570010) was added to all reactions in the dark. Vortexing was repeated before transferring reactions to a 96-well plate (200 µL per replicate well), which was incubated for 10 min at room temperature in the dark before measuring absorbance spectrophotometrically at 562 nm. To determine the ferrous iron chelating percentage of samples, the average blank absorbance was used for correction before subtracting sample absorbance from the average maximal signal control value, dividing by average maximal signal control value, and multiplying by 100. An EDTA standard curve was constructed and used to determine EDTA µM equivalents for each sample.

### 2.9 Antiviral Activity Assays

Because of consistent viral-related RNA-Seq pathway responses and past reports of agarikon as an effective antiviral treatment in both *in vitro* and *in vivo* contexts (9, 10, 12, 18), additional support for the antiviral activity of this agarikon strain became increasingly relevant. Extracts of agarikon mycelium have previously been evaluated for activity against an array of viruses as part of Project BioShield (12), including the strain used in the current study, which was tested as part of this program and follows methods reported in (12). In brief, EtOH extracts of agarikon mycelium were tested for antiviral activity in human cells against a wide range of viruses, including influenza A, using a neutral red assay and a visual cytopathic effect assay. Results are reported compared to the control antiviral drug ribavirin.

## 3 Results

### 3.1 Agarikon Mycelium Promotes Cellular Metabolic Activity and Proliferation Across Basal and LPS-Stimulated Conditions

To assess cytocompatibility and support selection of treatment concentrations, cell metabolic activity and proliferation were evaluated using MTT and fluorometric assays, respectively (Figure 1). The EtOH fraction of agarikon mycelium resulted in a consistent increase in cell viability, with the five lowest concentrations resulting in significant metabolic increases of at least 20% relative to the vehicle (Figure 1A), while an EtOAc fraction was observed to maintain viability, but not promote augmented metabolic outcomes (Figure 1B). Following treatment with the EtOH fraction cellular proliferation was maintained under both basal and LPS-stimulated conditions, with statistically significant increases compared to the vehicle observed across all tested concentrations (Figure 1C, 1D). These findings were supported by morphological observations of agarikon-treated cells (Supplementary Figure S1).

**Figure 1.**
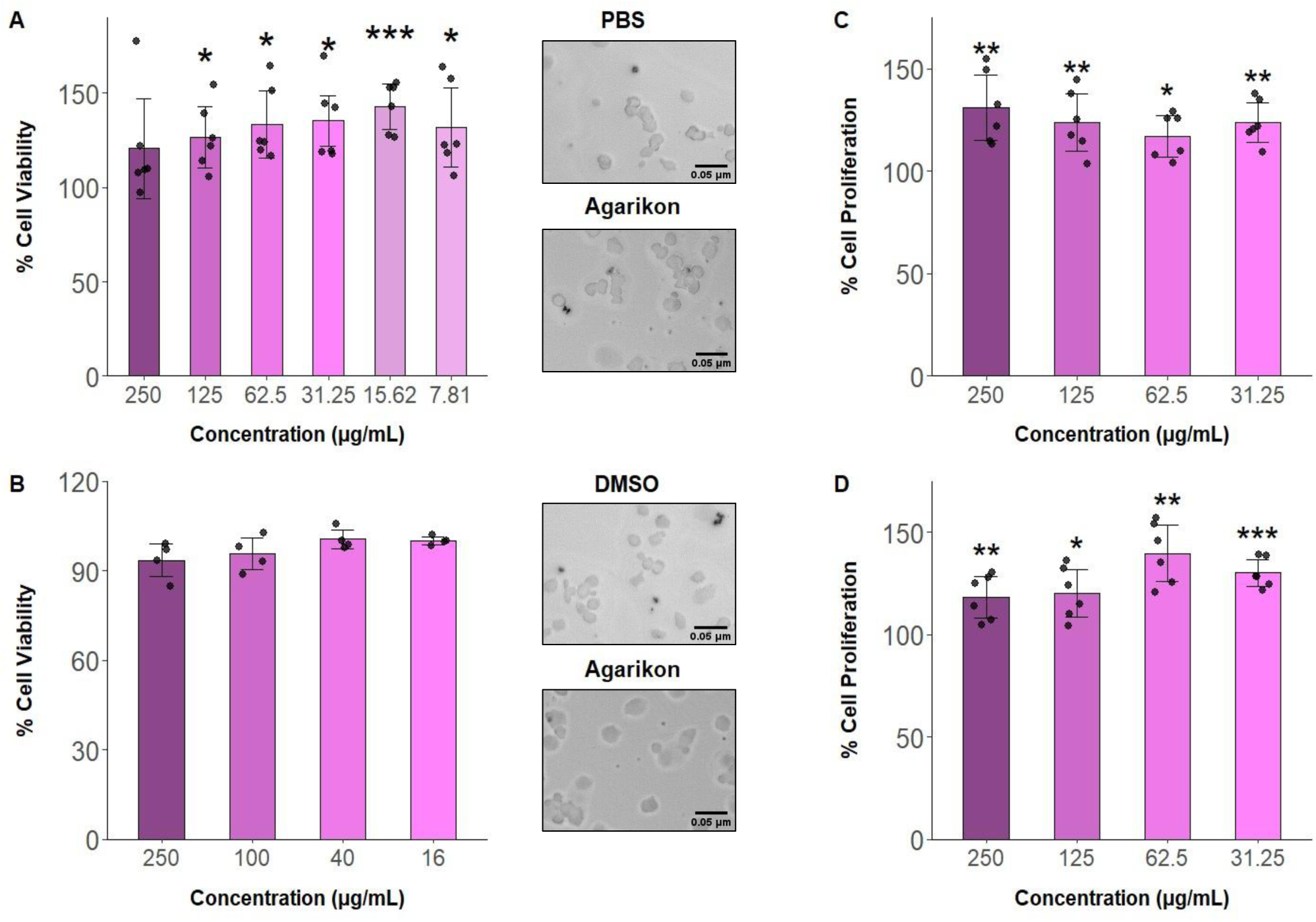
Percent cell viability of human peripheral blood mononuclear cells (PBMCs) treated with **(A)** hydroethanolic (EtOH) or **(B)** ethyl acetate (EtOAc) extracts of agarikon (*Fomitopsis officinalis*, syn*. Laricifomes officinalis*) mycelium, as evaluated by MTT assay. **(C)** Fluorometric assay of cell proliferation after treatment with an EtOH agarikon fraction relative to the vehicle (DMSO) under basal conditions and **(D)** LPS-stimulated conditions relative to vehicle with LPS. In all cases, n ≥ 4 wells per treatment concentration; significance evaluated relative to vehicle controls, two-tailed t-test, unequal variances, \**p* < 0.05; \*\**p* < 0.01; \*\*\**p* < 0.001.

### 3.2 Context-Dependent Transcriptomic Signatures of Immune and Stress Responses

PCA was performed for both basal and inflammatory RNA-Seq data to compare gene expression trends, revealing that agarikon treatment samples clustered separately from vehicle samples under both basal and inflammatory conditions (Figure 2A, 2B). PC1 under both basal (76.34%) and inflammatory (78.88%) conditions accounted for the majority of variance, while PC2 contributed minimally (8.22% and 5.58%, respectively). In both cases, the transcriptome was largely organized along a single dominant axis of variation, consistent with a strong treatment-associated transcriptional outcome.

**Figure 2.**
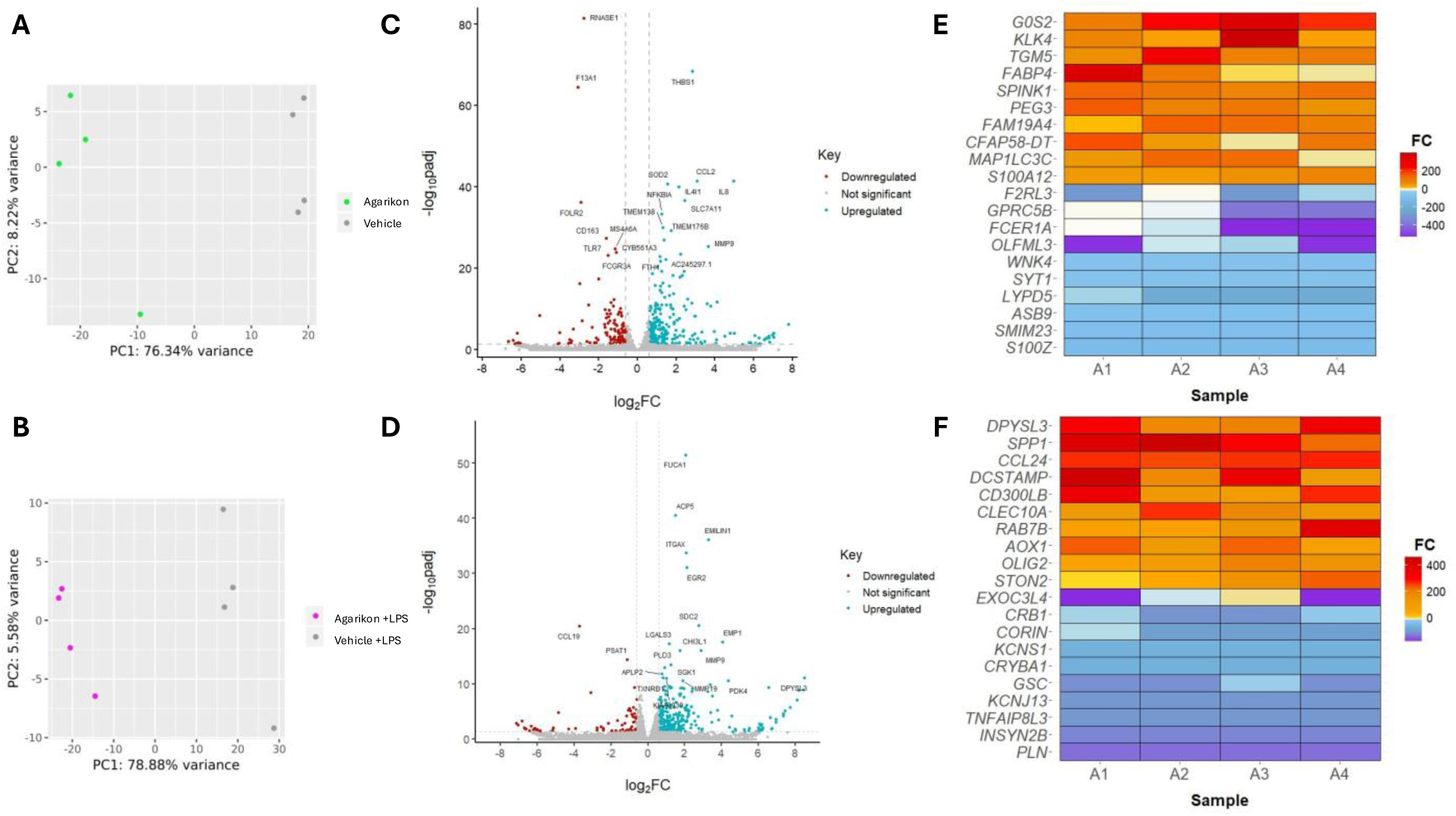
**(A)** Principal component analysis (PCA) for agarikon- and vehicle-treated samples under basal and **(B)** inflammatory conditions evaluating the similarities or differences between these treatment groups based on their transcriptomic clustering patterns (n = 4). **(C)** Volcano plot for agarikon-treated PBMCs relative to Vehicle under basal and **(D)** inflammatory conditions, displaying the magnitude of differential gene expression (log_2_FoldChange) in relation to statistical significance (-log_10_(*p*adj)). Statistically significant results are indicated in blue for downregulated genes and red for upregulated genes (n = 4, DESeq2, *p*adj < 0.05). **(E)** Top 10 up- and down-regulated genes by fold change (FC) for the agarikon treatment relative to Vehicle under basal and **(F)** inflammatory conditions (n = 4, DESeq2, *p*adj < 0.05).

Consistent with the clear treatment-associated separation observed by PCA, wide-ranging transcriptional differences were revealed between agarikon treatment relative to Vehicle (DESeq2 analysis, *p*adj < 0.05, Figure 2C). Under basal conditions, 351 genes were upregulated and 184 genes were downregulated. The most statistically significant results included ribonuclease A family member 1, pancreatic (*RNASE1*, *p*adj < 3.98 × 10^-82^, FC = -6.87), *CXCL8*/*IL8* (*p*adj < 3.89 × 10^-42^, FC = 31.02), superoxide dismutase 2 (*SOD2*, *p*adj < 1.75 × 10^-41^, FC = 2.93), interleukin 4 induced 1 (*IL4I1*, *p*adj < 1.07 × 10^-40^, FC = 4.37), and *NFKBIA* (*p*adj < 4.70 × 10^-34^, FC = 2.39). Highly upregulated transcripts included G0/G1 switch 2 (*G0S*, FC = 220.55), kallikrein related peptidase 4 (*KLK4*, FC = 130.18), and transglutaminase 5 (*FGM5*, FC = 126.47), while the most strongly downregulated transcripts included S100 calcium binding protein Z (*S100Z*, FC = -101.90), LY6/PLAUR domain containing 5 (*LYPD5*, FC = -74.56), and Fc fragment of IgE receptor Ia (*FCER1A*, FC = -16.50; Figure 2E).

Under inflammatory conditions, agarikon treatment upregulated 437 genes, representing a greater transcriptional activation response than observed under basal conditions, while a comparable number of genes were downregulated (181 genes). Among the most statistically significant results were alpha-L-fucosidase 1 (*FUCA1*, *p*adj < 2.99 × 10^-52^, FC = 4.12), C-C motif chemokine ligand 19 (*CCL19, p*adj < 3.76 × 10^-21^, FC = -13.07), galectin 3 (*LGALS3*, *p*adj < 5.04 × 10^-18^, FC = 2.24), and thioredoxin reductase 1 (*TXNRD1*, *p*adj < 7.77 × 10^-12^, FC = 1.75). Highly upregulated results included dihydropyrimidinase like 3 (*DPYSL3*, FC = 357.89), C-C motif chemokine ligand 24 (*CCL24*, FC = 292.25), and C-type lectin domain containing 10A (*CLEC10A*, FC = 173.84), whereas the most downregulated results included phospholamban (*PLN*, FC = -131.90), inhibitory synaptic factor family member 2B (*INSYN2B*, FC = -114.56), and TNF alpha induced protein 8 like 3 (*TNFAIP8L3*, FC = -100.04; Figure 2F).

Under basal conditions, significantly (*p*adj < 0.05) affected pathways in agarikon-treated PBMCs clustered around degranulation processes, TLR and interleukin immune signaling, and antioxidant/detoxification pathways (Figure 3A, 3C). After LPS stimulation, these pathways centered on appropriate response to simulated microbial invasion, including cytokine and TLR signaling and degranulation processes, while retaining signs of antioxidant and anti-inflammatory pathway engagement (Figure 3B, 3D). Individual transcript responses were consistent with these pathway results. Notable immune DEGs upregulated under basal conditions included *CXCL8/IL8, IL1RN/IL1RA,* IL-1 beta (*IL1B*)*, IL4I1, SOD2,* IL-18 binding protein (*IL18BP*), NF-κB subunit 2 (*NFKB2*), transcription factor RelB (*RELB*) and several cluster of differentiation (CD) markers, though *TLR4* and *TLR7* are significantly downregulated. While *IL8* transcript fold change (FC) values are robust, actual protein induction is much lower, with no agarikon treatment increasing IL-8 protein by more than 30 pg/mL (Supplementary Figure S2). Under inflammatory conditions*, IL8, IL1RA*, and *IL1B* were also upregulated, though not to the degree observed under basal conditions. In this challenged environment, while *TLR5, IL1A,* and many CD transcripts increased, they were attenuated in comparison to basal conditions, and opposite to basal conditions, *IL18* and the noncanonical NF-κB subunits (*NFKB2* and *RELB*) were downregulated (Figure 3E, 3F).

**Figure 3.**
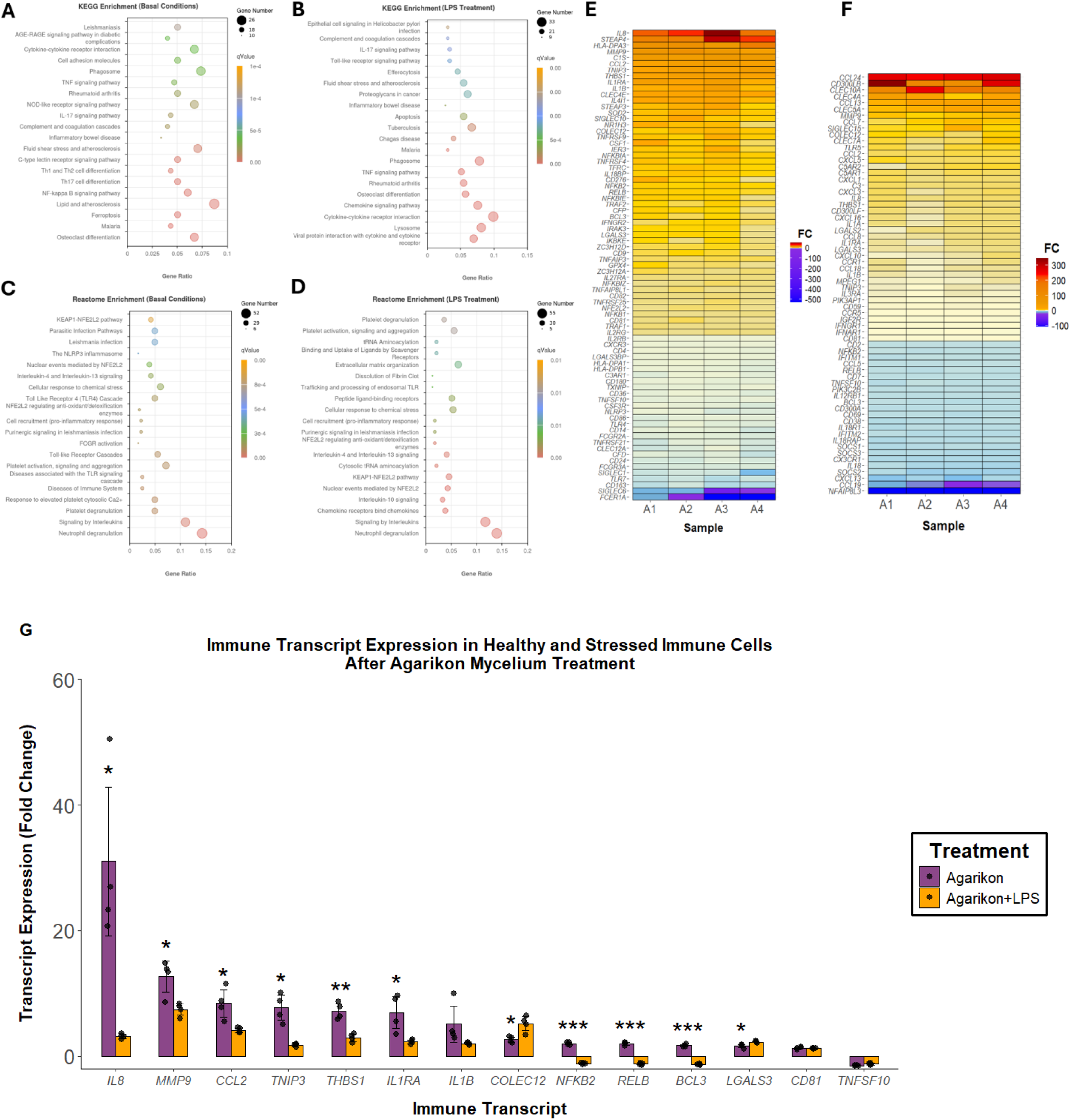
**(A)** KEGG dot plot of enriched pathways in agarikon-treated PBMCs relative to Vehicle under basal and **(B)** inflammatory conditions (n = 4, DESeq2, *p*adj < 0.05). **(C)** Reactome dot plot of enriched pathways in agarikon-treated PBMCs relative to Vehicle under basal and **(D)** inflammatory conditions (n = 4, DESeq2, *p*adj < 0.05). **(E)** Select immune-related differentially expressed genes (DEGs) identified in PBMCs treated with agarikon compared to Vehicle under basal and **(F)** inflammatory conditions (n = 4, DESeq2, *p*adj < 0.05). **(G)** Comparative levels of select immune DEGs shared by basal and inflammatory conditions (n = 4, significance testing performed by comparing basal condition averages to inflammatory condition averages, two-tailed t-test, unequal variances, \**p* < 0.05; \*\**p* < 0.01; \*\*\**p* < 0.001). Complete lists of immune DEGs under basal and LPS-inflammatory conditions can be found as Supplementary Tables S3 and S4.

When comparing FC values of immune DEGs in response to agarikon treatment under basal and inflammatory conditions, significant trends emerge (Figure 3G). Human PBMCs in LPS-induced inflammatory conditions produced ten-fold less *IL8* after treatment with agarikon compared to basal PBMCs, alongside a non-significant decrease in *IL1B* (*p* < 0.05). This is paired with a significant decrease in levels of *IL1RA*, as well as the active downregulation of *NFKB2* and *RELB*, key subunits of the noncanonical NF-κB pathway. Additionally, under basal conditions, the upregulation of immunoprotective factors such as *IL18BP* and *SOD2* is paired with the downregulation of key inflammatory actors such as *TLR4, TLR7*, and *CD14*. Agarikon treatment was associated with condition-dependent differential expression of TLR genes, including decreased *TLR4* and *TLR7* under basal conditions and increased *TLR5* under LPS-stimulated conditions, consistent with KEGG pathway annotations of upstream pattern recognition signaling components (Supplementary Figure S3). Overall, agarikon appears to induce context-dependent modulation of immune-associated gene expression under both basal and LPS-stimulated conditions.

### 3.3 Antioxidant Capacity of Agarikon Mycelium

Based on the transcript- and pathway-level antioxidant and ferroptosis results observed after agarikon treatment, DPPH radical scavenging activity (RSA) and ferrous iron chelating ability were also investigated (Figure 4). Agarikon mycelial extract exhibited significant, dose-dependent RSA in both water and ethanol extracts, and the response occurred rapidly, with >75% of total RSA achieved within 30 minutes. Activity was maintained across a broad concentration range, from 500 µg/mL to a minimum of 20.83 µg/mL (resuspended in water) and 18.5 µg/mL (resuspended in ethanol). Average RSA for this treatment was slightly higher when resuspended in ethanol, approaching 90% of the maximal control. Ferrous iron chelation activity also increased in a dose-dependent manner, with all tested concentrations displaying significantly more iron chelation activity than the maximal signal control (*p* < 0.05; Figure 4C). The magnitude of this response increased approximately four-fold across the concentration range tested before approaching a plateau at the two highest concentrations.

**Figure 4.**
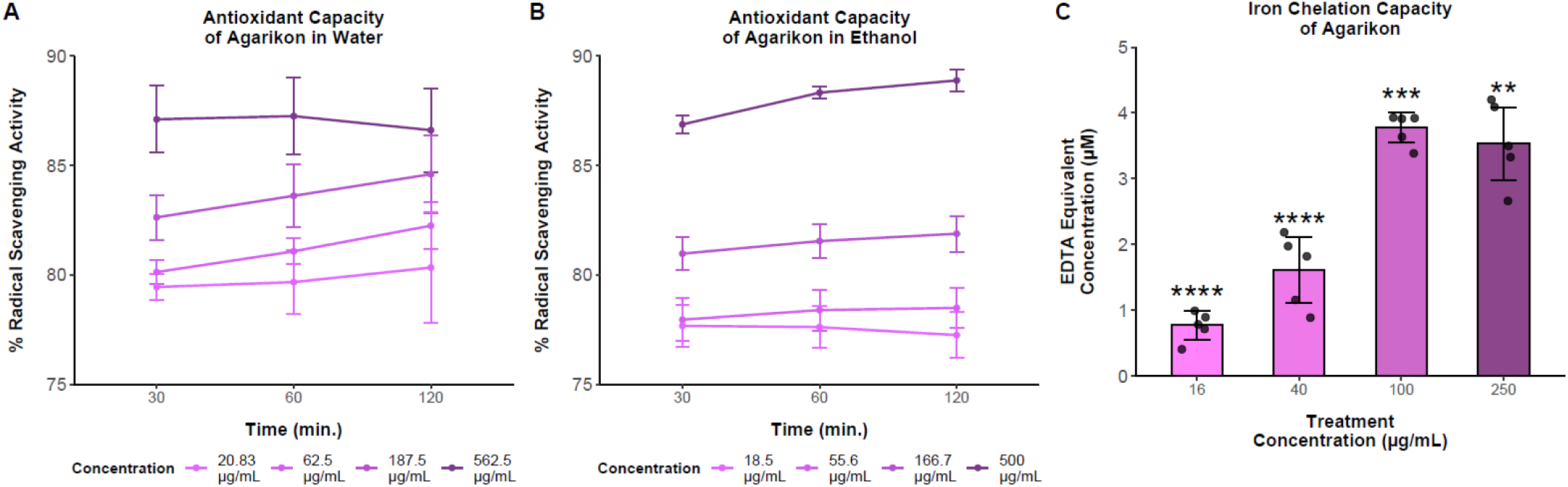
**(A)** Antioxidant capacity of agarikon over time in water and **(B)** ethanol as evaluated by DPPH assay (n = 7). **(C)** Ferrous iron chelation capacity of agarikon, expressed as EDTA concentration equivalents (n = 5; significance testing performed by comparing to maximum signal control, two-tailed t-test, unequal variances, \*\**p* < 0.01; \*\*\**p* < 0.001; \*\*\*\**p* < 0.0001).

### 3.4 Cytokine Modulation and Secondary Immune Signaling

To provide protein-level assessment of transcriptomic immune responses, ELISAs for select cytokine targets were also performed. Similar to transcript-level findings, the EtOH agarikon preparation induced significant increases in IL-1RA under basal conditions, while a similar induction was also observed for the EtOAc fraction (Figure 5A). Under inflammatory conditions, both the EtOH and EtOAc fractions produced small yet significant changes in IL-1α protein levels, with an increase observed for the EtOH fraction and a decrease observed for the EtOAc fraction (Figure 5B). Neither fraction resulted in substantial increases of IL-8 protein under basal conditions (<30 pg/mL; Supplementary Figure S2).

**Figure 5.**
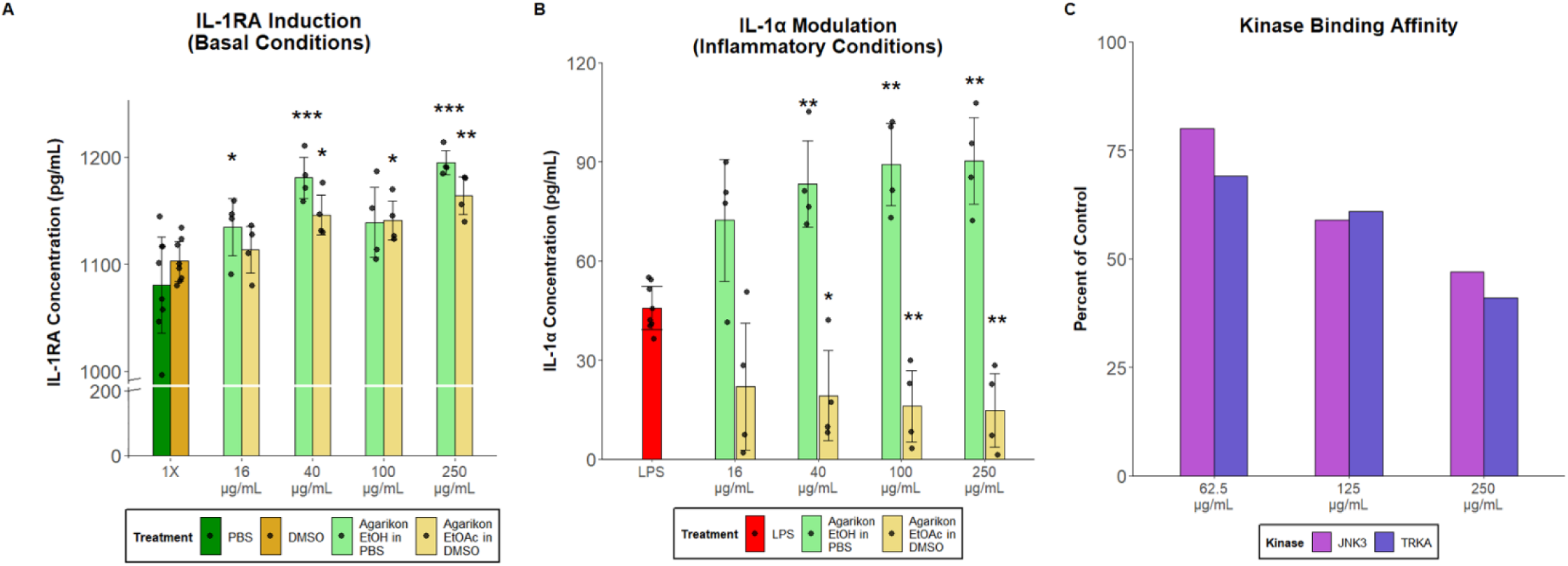
**(A)** IL-1RA cytokine responses in agarikon-treated human PBMCs under basal conditions (n = 4). **(B)** IL-1α responses under inflammatory conditions (n = 4). In all cases, significance was evaluated relative to respective vehicle controls (PBS, DMSO, or LPS) using two-tailed t-tests with unequal variances (\**p* < 0.05; \*\**p* < 0.01; \*\*\**p* < 0.001). **(C)** Kinase binding affinity of agarikon mycelium treatment toward JNK3 and TRKA as evaluated by Eurofins’ DiscoverX KINOMEscan profiling service. Lower values indicate greater engagement of the kinase by treatment and higher binding affinity overall.

MAPK binding affinity results (Figure 5C) indicate additional potential targets that may support the immune response observed in RNA-Seq and ELISA results. MAPK binding affinity screening focused on the EtOAc agarikon fraction, which is commonly used for enrichment of semi-polar secondary metabolites in target-based binding assays. These assays indicated moderate and concentration-dependent binding activity toward JNK3 and TRKA, with percent control values decreasing from approximately 75% to 50% across the concentration range, consistent with increased ligand displacement.

### 3.5 Influenza A Antiviral Activity

The antiviral activity of agarikon mycelium was of interest due to the consistently demonstrated antiviral activity of agarikon extracts, as well as the repeated presence of viral-related KEGG pathway results in the current study, including influenza A, Kaposi sarcoma-associated herpesvirus infection, tuberculosis, Epstein-Barr virus infection, hepatitis B, human T-cell leukemia virus 1 infection, and measles (*p*adj < 0.05). This specific strain of agarikon was previously included in a panel of polypore species extracts submitted for screening against numerous viruses as part of Project Bioshield, demonstrating especially notable activity against influenza A H3N2 (12; Figure 6). The selectivity index (SI) of a medicinal compound represents the ratio of its toxic concentration to its bioactive concentration, with values greater than 10 indicating a compound effective at a low concentration with a high concentration required for toxic effect, and values above 100 indicating notable therapeutic efficacy (29). As evaluated in neutral red and cytopathic effect assays, agarikon mycelium treatment demonstrated SI values of 260 or greater, up to 190% of those calculated for the antiviral drug ribavirin and far above the level accepted as both bioactive and safe.

**Figure 6.**
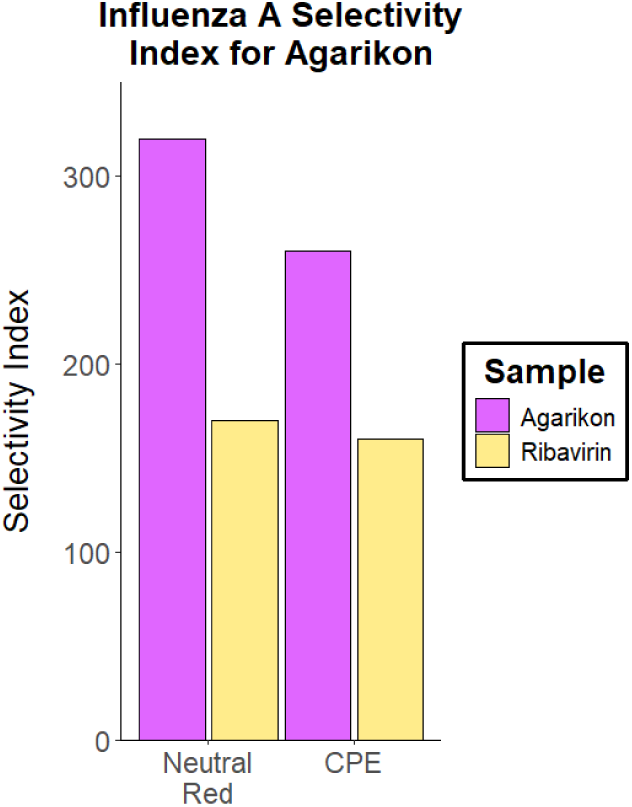
Selectivity index (SI) for agarikon mycelium against influenza A (H3N2) compared to ribavirin in neutral red and cytopathic effect (CPE) antiviral assays.

## 4 Discussion

### 4.1 Immune Response to Agarikon Treatment Under Basal Conditions

Under basal conditions, agarikon treatment of human PBMCs supported a balanced immune response characterized by concurrent engagement of immune-activating pathways alongside anti-inflammatory and antioxidant signatures, as well as antiviral and antimicrobial immune cascades. *TLR4* (FC = -1.68) and *TLR7* (FC = -2.85) were selectively downregulated alongside upregulation of other innate immune effectors, as reflected in KEGG and Reactome pathway enrichment (Figure 3A, 3C). Although fungal β-glucans are TLR agonists, the downregulation of TLR transcripts and the TLR co-receptor *CD14* (FC = -1.72) paired with upregulation of negative regulators such as IL-1 receptor-associated kinase 3 (*IRAK3*; FC = 1.64) suggests that this immune response is prompted by other fungal components (30, 31, 32, 33). Furthermore, this measured response includes the expression of a variety of complement cascade transcripts. While positive regulators such as complement component 1S (*C1S*; FC = 12.18) and properdin (*CFP*; FC = 1.71) are both engaged, the pathway is downregulated at many other stages of the complement cascade (complement C1q C chain (*C1QC*), FC = -1.42; complement C3a receptor 1 (*C3AR1*), FC = -1.43; complement factor D (*CFD*), FC = -1.99). Because *C3AR1* stimulus coactivates several TLRs, its downregulation may contribute to the notable *TLR4* and *TLR7* suppression observed (34, 35).

Agarikon treatment also resulted in modulation of cytokine signaling. *IL1RN*/*IL1RA* (FC = 6.94), which encodes IL-1RA protein, was even more upregulated than *IL1B* (FC = 5.09) and two separate agarikon preparations also induced significant IL-1RA protein production (Figure 5A). This pattern is consistent with regulatory modulation of the IL-1 axis rather than direct pro-inflammatory stimulation and aligns with immunomodulatory mechanisms proposed for agarikon and *T. versicolor* mycelium treatment in adjunctive COVID-19 vaccine contexts (8, 36). A disconnect between transcript and protein levels was observed for *IL8*, where robust transcriptional upregulation was not matched by comparable increases in secreted protein (Supplementary Figure S2). This pattern is consistent with a state of immune readiness or priming, in which cytokine-encoding transcripts are maintained at elevated levels without immediate translation into protein, enabling a more rapid response upon subsequent activation rather than reflecting active pro-inflammatory signaling at baseline (37, 38, 39). Similarly, IL-2 displayed divergent subunit expression (IL-2 receptor subunit gamma (*IL2RG*), FC = 1.28; IL-2 receptor subunit beta (*IL2RB*), FC = -1.20) paired with evidence of anti-inflammatory signals supported by IL-4 pathway modulation and significant *IL4I1* (FC = 4.37) upregulation (40). Taken together with T helper 2 cell (Th2) and IL-10-related pathway results, anti-inflammatory Th2 cell differentiation is likely, as IL-4 regulates the positive feedback loop for Th2 differentiation while also being produced by them, and IL-10 drives Th2 differentiation through regulation of dendritic cells (41). The strong engagement of these cascades and individual immune actors such as C-type lectin domain family 4 member E (*CLEC4E*/Mincle, FC = 4.97), which also positively regulates Th2 differentiation, supports a broader immunoregulatory profile associated with agarikon treatment (42, 43, 44). The noncanonical NF-κB pathway is known to be involved in positive regulation of anti-inflammatory Th2 cells (45), and agarikon treatment seems to modulate the NF-κB pathway, increasing transcript expression for subunits of both the canonical (*NFΚB1*, FC = 1.29) and noncanonical (*NFKB2*, FC = 1.99; *RELB*, FC = 1.94) NF-κB pathways, alongside several NF-κB inhibitors (*NFΚBIA*, FC = 2.39; NF-κB inhibitor epsilon (*NFΚBIE*), FC = 1.88; NF-κB inhibitor zeta (*NFΚBIZ*), FC = 1.39). These results are aligned with past reports of the anticancer effects of agarikon fruit body fractions driven by their upregulation of NF-κB proteins, including *NFKB1*/p50 and *NFKBIA*/IκBα, supporting the capacity of agarikon mycelium to provide a similar mechanistic drive towards immune modulation (13, 14, 15).

Transcripts engaged by agarikon mycelium treatment are largely involved with attenuation of inflammatory responses and redox modulation, either through direct antioxidant activity or negative regulation of inflammatory cellular markers. Antioxidant activity is reflected transcriptionally through the wide variety of antioxidant-related pathways and enzymes engaged such as the central regulator of mitochondrial redox balance *SOD2* (FC = 2.93), as well as glutathione peroxidase 4 (*GPX4*, FC = 1.46), peroxiredoxin 1 (*PRDX1*, FC = 1.36), thioredoxin (*TXN*, FC = 2.25), *TXNRD1* (FC = 1.39), and antioxidant 1 copper chaperone (*ATOX1*, FC = 1.85; 4, 13, 47). Thioredoxin-interacting protein (*TXNIP*, FC = -1.44), the *TXN* inhibitor, is also downregulated. Additionally, several iron recycling pathway components are notably upregulated (six-transmembrane epithelial antigen of the prostate 4 (*STEAP4*), FC = 27.35; *STEAP3*, FC = 3.06; transferrin receptor (*TFRC*), FC = 2.26; cytochrome B561 family member A3 (*CYB561A3*), FC = 2.60) alongside family with sequence similarity 20, member C (*FAM20C*, FC = 2.28), which helps maintain ER redox homeostasis (47, 48, 49). Upregulation of these transcripts is likely responsible for the ferroptosis pathway response observed (Figure 3A, 3C), due to its reactive oxygen species (ROS)-based mechanism of action (50). This is further supported by the lack of cytotoxicity observed across a range of agarikon treatment concentrations and preparations, in conjunction with demonstration of its antioxidant ability and iron chelation potential (Figures 1 and 4). Although the functional consequences of agarikon binding to JNK3 remain to be determined, this interaction (Figure 5C) may be relevant to the coordinated redox remodeling observed in the present study, as JNK signaling has been implicated in oxidative stress responses and ferroptosis-associated cell death (51, 52).

Agarikon mycelium influenced a range of innate immune pathways spanning myeloid, dendritic, and cytokine signaling. For example, leukocyte immunoglobulin-like receptor subfamily B member 4 (*LILRB4*, FC = 1.77) helps inhibit myeloid cytokine production, while phospholipase A2 group VII (*PLA2G7*, FC = 3.62) inactivates inflammatory signaling molecules and decreases monocyte activity (53, 54, 55), similarly to *IL18BP* (FC = 2.25), which inhibits inflammatory IL-18 and the downstream production of IFN-γ (56). Secreted protein acidic and cysteine rich (*SPARC*, FC = 5.18) interacts with thrombospondin-1 (*THBS1*, FC = 7.17), which then regulates activity of a wide array of ligands; while the THBS1 interactome is complex, it is known to negatively regulate dendritic activation and can play an anti-inflammatory role (57, 58). Notably, transmembrane protein 176A (TMEM176A, FC = 3.16) is associated with immature dendritic cells, a hallmark of active immunomodulation or a lack of inflammatory signals as immature dendritic cells release much lower levels of inflammatory cytokines and T cell activators compared to mature cells (59, 60).

Overall, while immunoactivating transcripts are upregulated by agarikon mycelial treatment under basal conditions, the accompanying engagement of regulatory and anti-inflammatory actors seems to pair with its demonstrated antioxidant activity to buffer the immune system away from extreme responses while at rest.

### 4.2 Immune Modulation of Agarikon Treatment After LPS Stimulation

Overall, the general pattern of immune engagement alongside simultaneous anti-inflammatory action was also observed under inflammatory conditions, though with additional (and appropriate) stimulation of immune effectors in response to LPS challenge. First, pathway results repeatedly implicate cytokine and interleukin signaling, and key messengers (*IL1A*, FC = 2.56; *IL1B*, FC = 2.01; *IL8*, FC = 3.14; *TLR5*, FC = 4.28) are significantly upregulated, indicating an appropriate response to LPS stimulation, where robust but regulated activation of these pathways is required for effective immune function. Interestingly, differences in extraction method resulted in a divergent IL-1α protein response, where the EtOH agarikon fraction resulted in small (<50 pg/mL) but significant increases in IL-1α, while the EtOAc fraction resulted in similarly significant decreases in IL-1α protein levels (Figure 5B). Overall, the pg/mL levels of IL-1α protein induced under inflammatory conditions are two orders of magnitude lower than the amount of IL-1RA induced under basal conditions, which aligns with the 5-to 100-fold greater concentrations of IL-1RA required to antagonize IL-1α activity (61, 62).

Cytokine transcript upregulation is paired with the downregulation of three suppressor of cytokine signaling transcripts (*SOCS1*, FC = -1.53; *SOCS2*, FC = -2.18, *SOCS3*, FC = -1.65), key regulators of cytokine signaling feedback inhibition (63, 64). The inflammatory IL-18 axis is also suppressed, with downregulation of several IL-18 receptor components (IL-18 receptor 1 (*IL18R*), FC = -1.45; IL-18 receptor accessory protein (*IL18RAP*), FC = -1.51) and *IL18* (FC = -1.84) itself (65). This early response is also associated with the canonical NF-κB pathway, as it is positively regulated by TLR5 and IL-8 signaling (66). Additionally, both subunits of the noncanonical NF-κB pathway (*RELB*, FC = -1.28; *NFKB2*, FC = -1.19) are moderately repressed, while C-type lectin domain family 7 member A (*CLEC7A*/Dectin-1, FC = 4.71), a fungal β-glucan receptor, only appears as an upregulated transcript under inflammatory conditions. Dectin-1 is a well-established activator of the canonical NF-κB pathway through complexing with TLRs, which aligns with past observations of TLR and NF-κB engagement after agarikon treatment of cancer cells (13, 14, 15, 67, 68, 69, 70). Notably, TLR and NF-κB subunit transcript expression exhibited opposing patterns under basal and inflammatory conditions, with reduced *TLR* and increased NF-κB subunit transcript expression at baseline reversing following LPS stimulation. Together, these findings suggest a context-dependent immunomodulatory response.

In contrast to the basal response, agarikon treatment under LPS stimulation seems to have prompted complement cascade activation (complement C3 (*C3*), FC = 3.21; complement C5a receptor 1 (*C5AR1*), FC = 3.59; complement C5a receptor 2 (*C5AR2*), FC = 3.81), indicating an enhanced engagement of innate immune effector mechanisms. However, the C-type lectins expressed reflected a balance between immune activation and immune regulation. While there was notable upregulation of *CLEC10A* (FC = 173.84), which promotes IL-10 production by dendritic cells, its induction was coupled with *CLEC5A* (FC = 10.98), a myeloid innate immune checkpoint that participates in pro-inflammatory signaling (71, 72). Lectin modulation was also evident among galectins involved in innate and adaptive immune regulation, including galectin-2 (*LGALS2*, FC = 2.46) and *LGALS3* (FC = 2.24). *LGALS3* has complex signaling effects, helping to upregulate inflammatory signals via the NLR family pyrin domain containing 3 (NLRP3) inflammasome, IL-1β production, and neutrophil recruitment, while also preventing over-engagement by raising the threshold for T cell activation and positively regulating anti-inflammatory M2 macrophages (73, 74, 75, 76, 77, 78, 79). This may suggest a role for galectin signaling in agarikon-mediated modulation of the IL-1 axis.

While a robust immune response to challenge is desirable, excessive inflammation can be detrimental. Agarikon treatment appears to maintain this signaling balance through influencing a wide variety of immune signaling networks to aid cellular response to LPS. For example, macrophage-expressed 1 (*MPEG1*, FC = 1.79) is an antimicrobial pore-forming protein upregulated by LPS, and stabilin-1 (*STAB1*, FC = 9.45) is a scavenger receptor that helps to clear LPS from the bloodstream (80, 81). This response was also characterized by modest downregulation of seven aminoacyl-tRNA synthetase transcripts (ARSs), including multiple cytosolic isoforms, which play roles beyond protein translation in inflammatory and oncogenic signaling (82, 83, 84). A small set of transcripts associated with inflammatory activation and poor cancer prognosis, including interferon induced transmembrane protein 1 (*IFITM1*, FC = −1.21), adhesion G protein-coupled receptor E5 (*ADGRE5*, FC = −1.30), and *CCL19* (FC = −13.07), were also reduced, consistent with attenuation of inflammatory amplification processes (85, 86, 87, 88). Conversely, expression of *FUCA1* (FC = 4.12) is associated with better cancer survival rates and may play a direct suppressive role in tandem with galactose-3-O-sulfotransferase 4 (*GAL3ST4*, FC = 2.23), which promotes macrophage M2 transition (89, 90, 91). Additionally, transmembrane protein 119 (*TMEM119*, FC = 10.33) helps suppress microglial inflammatory reactions after strokes (92), and this strongly expressed candidate is notable in the context of agarikon’s demonstrated binding affinity for TRKA (Figure 5C), the primary receptor of nerve growth factor (NGF) in the brain.

Finally, agarikon-induced redox remodeling was maintained under LPS stimulation, indicating that antioxidant- and ferroptosis-related protective signatures persist despite inflammatory ROS generation. Pathway-level signals point to engagement of iron recycling and ROS detoxification processes linked to resolution of inflammatory oxidative stress, supporting maintenance of redox homeostasis during immune activation. This response was supported by the upregulation of glutathione-disulfide reductase (*GSR*, FC = 1.45), *TXN* (FC = 1.71), and *TXNRD1* (FC = 1.75), alongside ferroptosis suppressor solute carrier family 7 member 11 (*SLC7A11*, FC = 3.20) and NAD(P)H quinone dehydrogenase 1 (*NQO1*, FC = 2.29), a stress-inducible enzyme involved in cellular protection against redox imbalance (93, 94, 95). Overall, the immune engagement prompted by agarikon treatment of human PBMCs after LPS challenge seems to be tightly regulated by concurrent immune-pacifying and immune-engaging signals, with parallel maintenance of redox balance.

### 4.3 Translational Relevance of Agarikon Immunomodulation

The transcriptomic immunomodulatory profile showed a clear overlap with antiviral-associated pathways, supporting a potential mechanistic basis for the well-established antiviral activity of agarikon observed in functional and clinical contexts (9, 10, 12, 18). Agarikon treatment engaged key mediators known to support innate immune activation and viral clearance, such as *IL8* and *IL1B* (Figures 3G and 7A, 7B), while also inducing significant virus-related pathway results (Figure 3A, 3B, 3C, 3D), notably including influenza A (96, 97, 98). These system-level insights align with functional antiviral profiling of agarikon mycelial extract against influenza A (H3N2) infection (Figure 6), which demonstrated SI values exceeding 300 in some assays and consistently higher activity than ribavirin alongside low cytotoxicity. Influenza-related antiviral activity of agarikon has also been reported in independent studies, including a prior investigation of the antiviral activity of aqueous extracts of agarikon mycelium against H5N1 and H3N3, which produced significant reductions in viral titers at concentrations that remained non-cytotoxic (18). Furthermore, the strain of agarikon mycelium employed in the current study has been evaluated clinically in combination with *T. versicolor* mycelium in multiple contexts, using unfermented rice as a placebo control. One clinical trial (9) investigating the use of this fungal preparation as a COVID-19 treatment has established that this agarikon and *T. versicolor* mycelium blend is a safe and well-tolerated intervention associated with reduced number and severity of COVID-19 symptoms, with supportive preclinical findings demonstrating significant reductions in viral entry in mammalian epithelial cells (Vero-TMPRSS2). Agarikon and *T. versicolor* mycelium also show promise as a safe COVID-19 vaccine adjunct, where the blend was associated with reduced vaccine-related side effects without compromising SARS-CoV-2 antibody responses, and potential indications of increased antibody levels for at least six months in patients without prior COVID-19 exposure (8).

**Figure 7.**
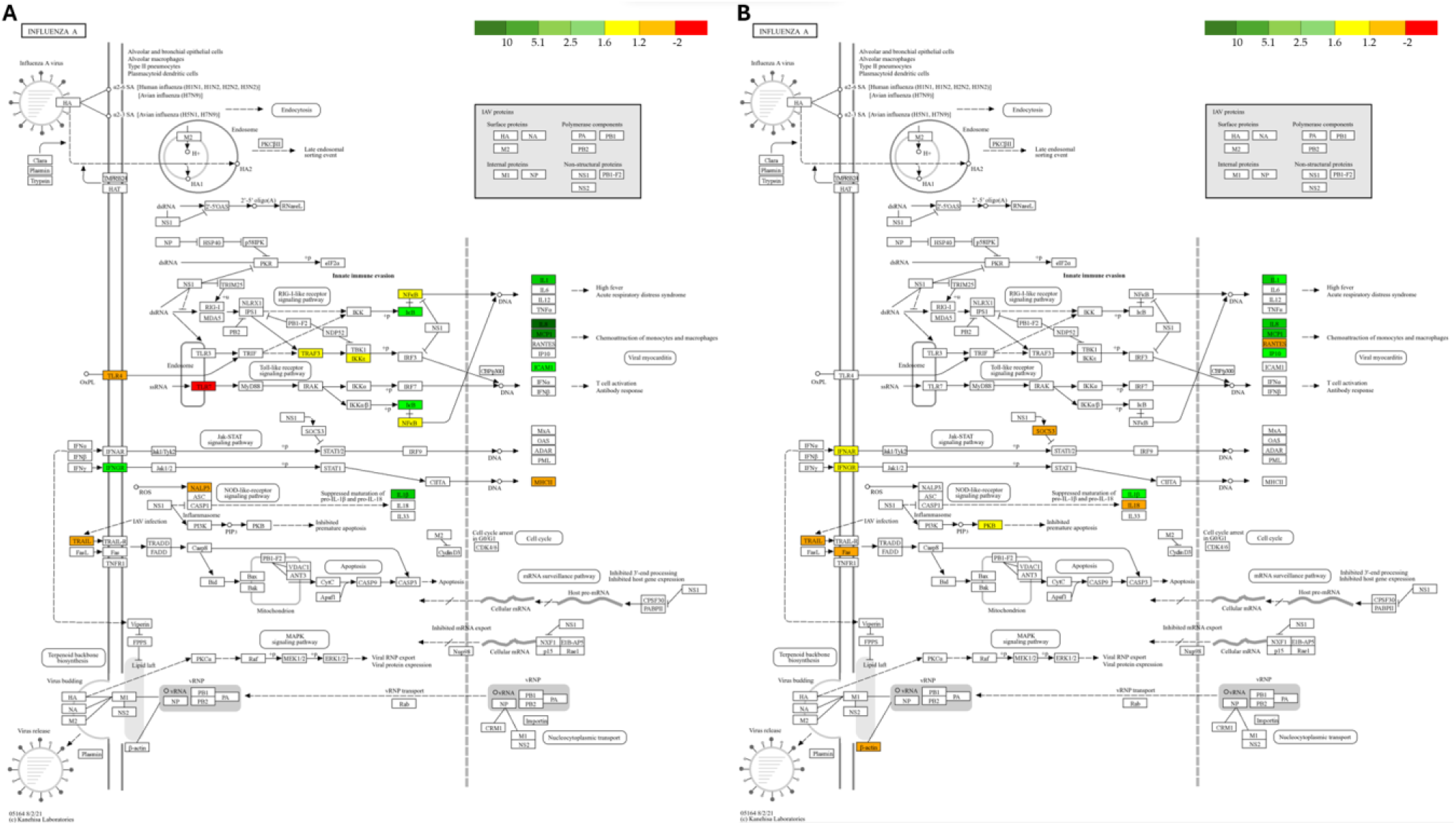
KEGG pathway for influenza A with genes affected by agarikon treatment under **(A)** basal and **(B)** inflammatory conditions highlighted according to average FC (n = 4, DESeq2, *p*adj < 0.05).

In addition to its inclusion in traditional European medicinal preparations termed “elixir[s] of long life,” modern research suggests a potential role for agarikon in supporting longevity-related biological processes, particularly under stress conditions (99, 100). The same properties that contribute to the antiviral and antimicrobial action of agarikon raise the possibility that this species could ameliorate age-related diseases (ARDs) driven by long-term inflammation (2, 4, 6, 10, 12, 14, 15). These findings may also have relevance to healthy ageing, as immune dysregulation and redox imbalance are central features of ARDs and inflammaging, the long-term, low-grade inflammation induced by chronic stimulation of the innate immune system (101, 102, 103, 104). The coordinated antioxidant and context-dependent immunomodulatory responses observed here are consistent with mechanisms associated with resilience to chronic inflammatory stress, although these implications would benefit from clinical validation (105, 106, 107, 108, 109). The antioxidant profile supported by agarikon aligns with findings from a D-galactose-induced mouse ageing model, in which agarikon improved antioxidant enzyme activities (SOD, CAT, and GSH-Px), reduced malondialdehyde (MDA) levels, and improved tissue indices associated with ageing (100). Collectively, these findings support a role for agarikon mycelium in regulating host responses across multiple stress contexts, characterized by immune buffering and redox homeostasis.

## 5 Conclusion

Despite millennia of traditional use and modern clinical application, agarikon treatment remains incompletely characterized with respect to mechanistic immune outcomes. Building on findings reported for the fruit body, this work highlights coordinated immunoregulatory effects elicited by agarikon mycelium, providing context for preclinical studies and clinical investigations of individual and blended preparations including agarikon and supporting hypotheses regarding its specifics of action, such as IL-1 axis modulation. While mycelial and fruit body preparations seem to produce both overlapping and distinct biological effects, this study further supports the immune-buffering capacity of mycelium-based treatment and highlights overlap with fruit body findings, including engagement of NF-κB signaling. Given the endangered status of agarikon, mycelial cultivation offers a scalable and sustainable approach to accessing its bioactive potential, underscoring the importance of continued research on this material. Overall, these findings support a role for agarikon in promoting adaptive resilience under stressed conditions, characterized by coordinated immunomodulatory activity and maintenance of redox balance.

## 6 Data Availability Statement

Differential gene expression results are provided in Table S1 and S2. All data from this manuscript are available upon reasonable request to the corresponding author.

## 7 Author Contributions

Conceptualization, C.B. and E.D.; Methodology, C.B., E.D. and J.K.; Software, E.D. and Z.J.B.; Validation, E.D. and J.K.; Formal Analysis, E.D.; Investigation, J.K. and E.D.; Data Curation, E.D. and J.K.; Writing—Original Draft Preparation, E.D. and C.B.; Writing—Review & Editing, E.D., C.B., J.K., Z.J.B., and P.S.; Visualization, E.D., Z.J.B. and C.B.; Supervision, C.B.; Project Administration, C.B. and E.D. All authors have read and agreed to the published version of the manuscript.

## 9.#Acknowledgments

We gratefully acknowledge Kyle Meyer, Patrick Bennett, and Jacqueline Morgado for their valuable insight and support.

## **11** Supplementary Material

**Table S1:** Complete DEG list with unfiltered RNA-Seq results for the transcriptomic profiles of human PBMCs treated with agarikon mycelium compared to the PBS vehicle under basal conditions.

**Table S2:** Complete DEG list with unfiltered RNA-Seq results for the transcriptomic profiles of human PBMCs treated with agarikon mycelium compared to the LPS vehicle under LPS inflammatory conditions.

**Table S3:** Immune-related DEGs elicited by agarikon mycelium treatment in PBMCs under basal conditions relative to the PBS vehicle.

**Table S4:** Immune-related DEGs elicited by agarikon mycelium treatment in PBMCs under inflammatory conditions relative to the LPS vehicle.

**Figure S1:** Representative images of agarikon- and vehicle-treated human immune cells (PBMCs) used for fluorometric cell proliferation assays under basal and LPS-induced inflammatory conditions.

**Figure S2**: IL-8 cytokine responses in agarikon-treated human PBMCs under basal conditions (n = 4). Significance was evaluated relative to respective vehicle controls (PBS or DMSO) using two-tailed t-tests with unequal variances (\**p* < 0.05; \*\**p* < 0.01; \*\*\**p* < 0.001).

**Figure S3: (A)** TLR transcripts affected by agarikon treatment under basal and inflammatory conditions (n = 4, DESeq2, all TLR responses *p*adj < 0.05 included). **(B)** KEGG pathway for the toll-like receptor signaling pathway with genes affected by agarikon treatment in human PBMCs under basal and **(C)** inflammatory conditions highlighted according to average FC (n = 4, DESeq2, *p*adj < 0.05).

