## Supplementary Figures S1, S2, S3 for "Adaptive Resilience: Agarikon Mycelium Modulates Immune Responses and Provides Oxidative Stress Buffering in Human Immune Cells"

### *Supplementary Material*

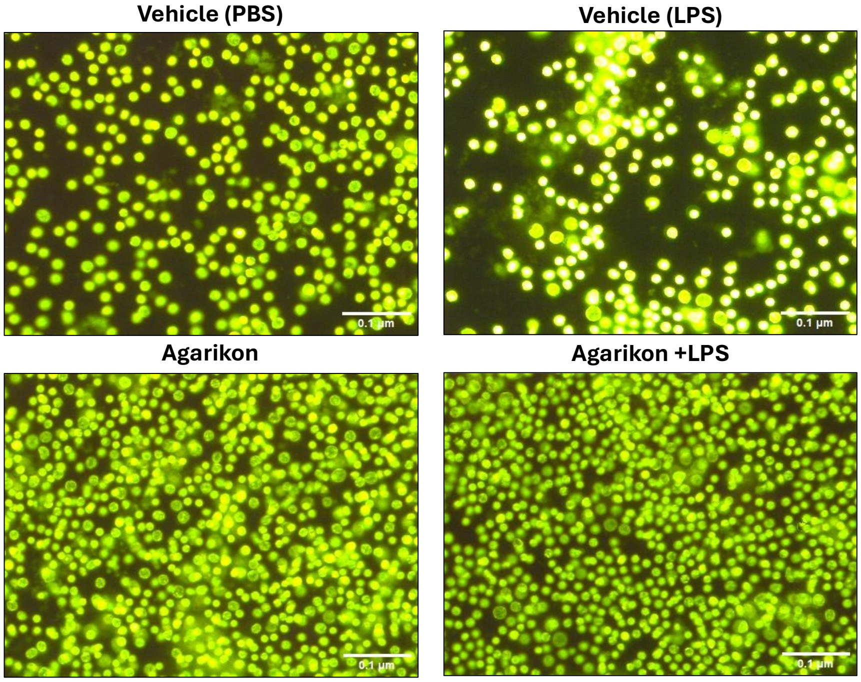


**Supplementary Figure 1.** Representative images of agarikon- and vehicle-treated human immune cells (PBMCs) used for fluorometric cell proliferation assays under basal and LPS-induced inflammatory conditions.

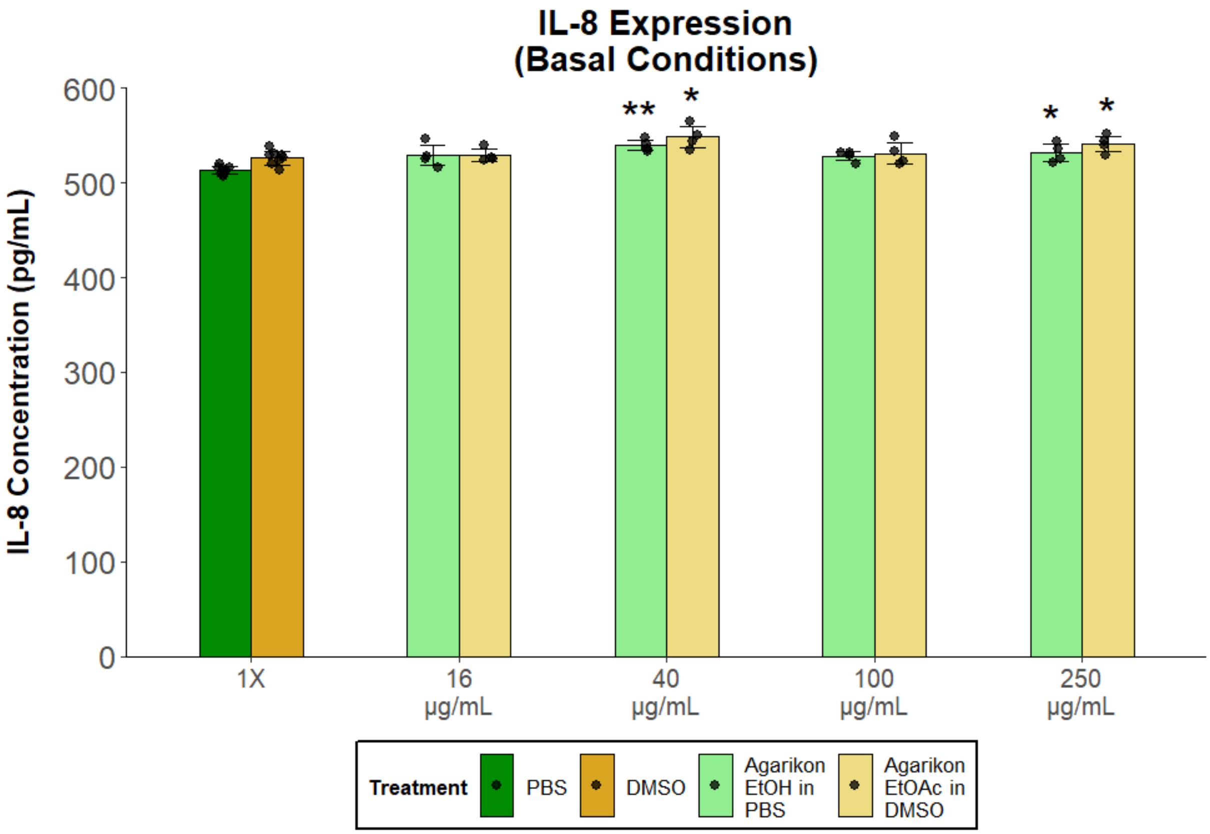


**Supplementary Figure 2.** IL-8 cytokine responses in agarikon-treated human PBMCs under basal conditions (n = 4). Significance was evaluated relative to respective vehicle controls (PBS or DMSO) using two-tailed t-tests with unequal variances (*$p < 0.05$; **$p < 0.01$).

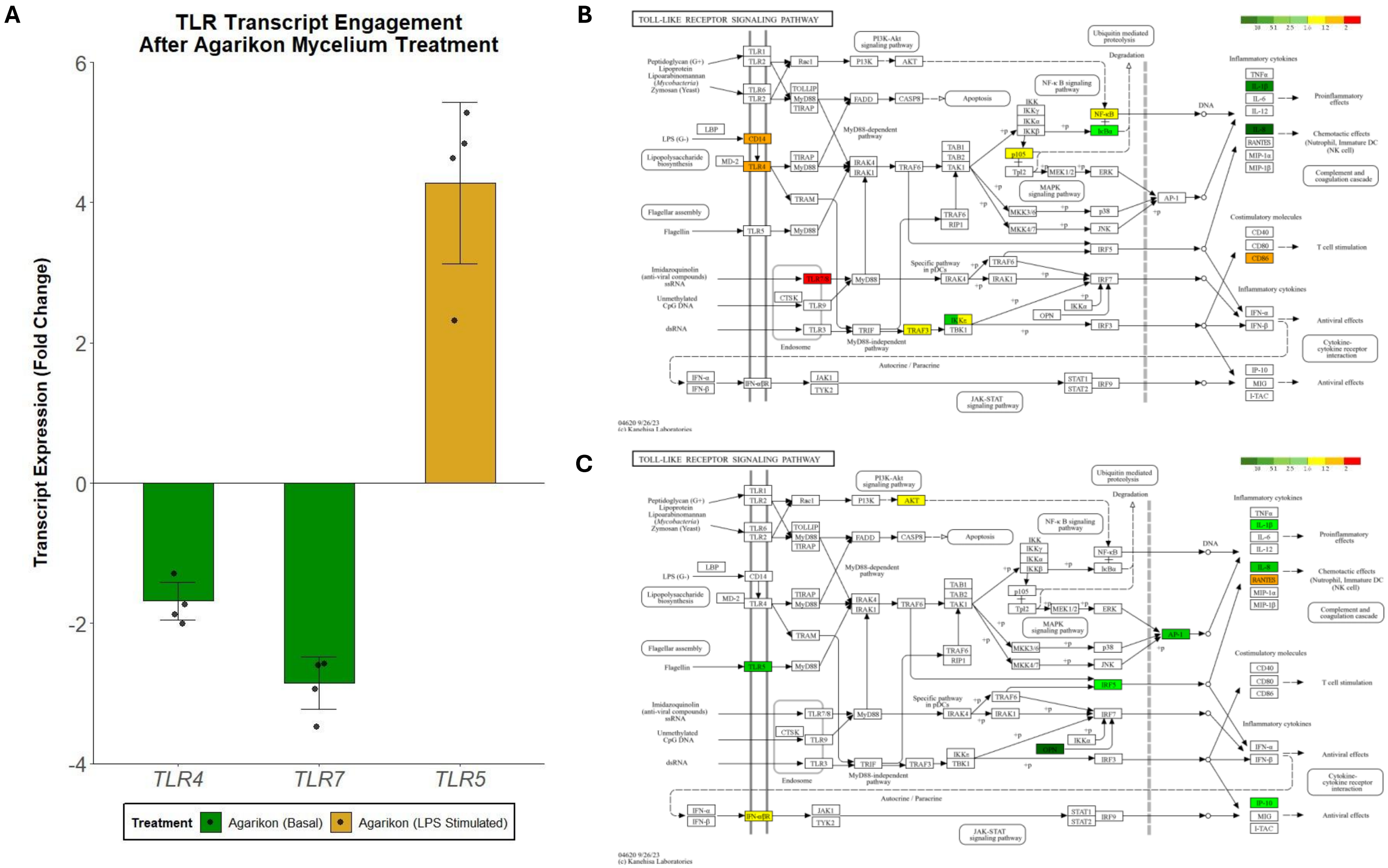


**Supplementary Figure 3. (A)** Toll-like receptor (TLR) transcripts affected by agarikon treatment under basal and inflammatory conditions (n = 4, DESeq2, all TLR responses *p*adj < 0.05 included). **(B)** KEGG pathway for the TLR signaling pathway with genes affected by agarikon treatment in human PBMCs under basal and **(C)** inflammatory conditions highlighted according to average fold change (n = 4, DESeq2, *p*adj < 0.05).
